# Scaling computational genomics to millions of individuals with GPUs

**DOI:** 10.1101/470138

**Authors:** Amaro Taylor-Weiner, François Aguet, Nicholas J. Haradhvala, Sager Gosai, Shankara Anand, Jaegil Kim, Kristin Ardlie, Eliezer M. Van Allen, Gad Getz

**Author notes:** Address correspondence to: Gad Getz. these authors contributed equally (Amaro Taylor-Weiner, François Aguet).

## Abstract

Current genomics methods were designed to handle tens to thousands of samples, but will soon need to scale to millions to keep up with the pace of data and hypothesis generation in biomedical science. Moreover, costs associated with processing these growing datasets will become prohibitive without improving the computational efficiency and scalability of methods. Here, we show that recently developed machine-learning libraries (TensorFlow and PyTorch) facilitate implementation of genomics methods for GPUs and significantly accelerate computations. To demonstrate this, we re-implemented methods for two commonly performed computational genomics tasks: QTL mapping and Bayesian non-negative matrix factorization. Our implementations ran > 200 times faster than current CPU-based versions, and these analyses are ∼5-10 fold cheaper on GPUs due to the vastly shorter runtimes. We anticipate that the accessibility of these libraries, and the improvements in run-time will lead to a transition to GPU-based implementations for a wide range of computational genomics methods.

## Background

Current methodologies for analyzing genomic data were designed for datasets with tens to thousands of samples, but due to the continuing drop in sequencing costs and the growth in large-scale genomic projects, data sizes are reaching millions of samples or individual cells. The need for increased computational resources, most notably runtime, to process these growing datasets will become prohibitive without improving the computational efficiency and scalability of methods. For example, methods in population genetics, such as genome-wide association studies (GWAS) or mapping of quantitative trait loci (QTL), involve billions of regressions between genotypes and phenotypes. Currently, the state-of-the-art infrastructures for performing these tasks are large-scale clusters of central processing units (CPUs), often with thousands of cores, resulting in significant costs^1^ (960 cores on a standard Google Cloud machine currently costs $7,660.80 per day of compute). In contrast to CPUs, a single graphical processing unit (GPU) contains thousands of cores at a much lower price per core (Nvidia’s P100 has 3,584 cores and currently costs $35.04 per day of compute).

Previous work has already demonstrated the benefits of using GPUs to scale bioinformatics methods^2–4^. However, these implementations were often complex and based on specialized libraries, limiting their extendibility and adoption. In contrast, recently developed open-source libraries such as TensorFlow^5^ or Pytorch^6^ make development of GPU-compatible tools widely accessible to the research community. We hypothesized that the use of these libraries would result in significant improvements in computational efficiency and enable scaling computational genomics methods to millions of samples.

## Results and Discussion

To study the efficiency and benchmark the use of TensorFlow and PyTorch for large-scale genomic analyses on GPUs, we re-implemented methods for two commonly performed computational genomics tasks: (i) QTL mapping^7, 8^ (which we call tensorQTL) and Bayesian non-negative matrix factorization^9^ (named SignatureAnalyzer-GPU). We executed the same scripts in identical environments (configured with and without a GPU) and also compared them to previous CPU-based implementations. For SignatureAnalyzer-GPU (SA-GPU), we used the mutation counts matrix generated by the Pan-Cancer Analysis of Whole Genomes (PCAWG) Consortium, which contains 2,624 tumors represented by 1697 mutational features of somatic single nucleotide variants and short insertions and deletions (defined based on their sequence contexts)^10^. Our PyTorch implementation ran approximately 200 times faster on a GPU than the current implementation of SignatureAnalyzer (SA) in R, with mean times for 10,000 iterations of 1.09 minutes using SA-GPU vs. 194.8 minutes using SA (Fig. 1a). Using simulated data, we showed that SA-GPU scales linearly with the number of samples (Supplementary Fig. 1a). When applied to previously published mutational signatures generated by SA^11^, we found the results of the two methods were essentially identical taking into account the stochastic nature of the underlying algorithm (mean R^2^ = 0.994, min R^2^ = 0.96; Fig. 1b).

**Figure 1:**
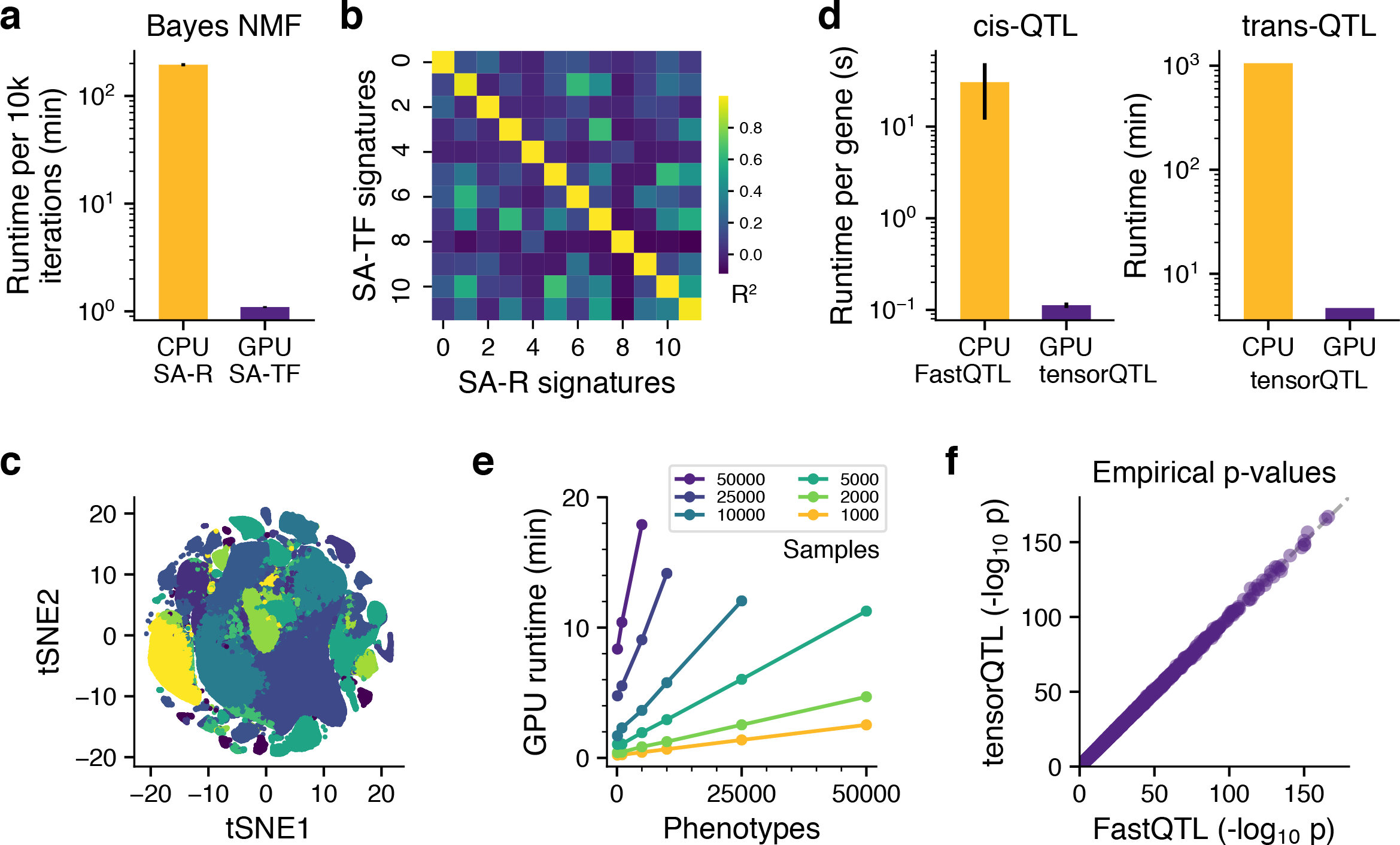
Performance of GPU implementations for QTL mapping and signature analysis. a) Average run time to compute 10,000 iterations of Bayes NMF using SignatureAnalyzer (SA) in R (purple) and SignatureAnalyzer-GPU (SA-GPU; gold). b) Correlation heat map of mutation signatures derived from the R and GPU implementations of SignatureAnalyzer using the same input mutation counts matrix. c) t-Distributed Stochastic Neighbor Embedding (t-SNE) of 1 million embryonic mouse brain cells. Colors indicate clustering based on SA-GPU decomposition performed in ∼15 minutes. d) Comparison of runtimes for *cis*-QTL (FastQTL on CPU; gold and tensorQTL on GPU; purple) and *trans*-QTL (tensorQTL on CPU and GPU). e) GPU runtime of tensorQTL for the indicated numbers of samples and phenotypes. f) Empirical *cis*-eQTL p-values from the V7 GTEx release replicated using tensorQTL. Error bars indicate standard deviation of the mean.

To further demonstrate the scalability of the Bayesian non-negative matrix factorization to millions of data points, we used SA-GPU to identify cell types and their associated transcriptional programs from single cell RNA sequencing of 1 million mouse brain cells (SRA: SRP096558, Figure 1C). The average time per SA-GPU run was 14.5 minutes (using V100 Nvidia GPU; n = 10) corresponding to an average of 6,853 matrix updates per run. A similar analysis on a CPU would require >2 days per run. Our analysis was able to identify 32 distinct transcriptional programs (Figure 1C).

For tensorQTL benchmarking, we generated random data representing up to 50,000 people, each with 10^7^ genotypes representing common variants. For each individual, we also simulated up to 50,000 phenotypes, resulting in 500 × 10^9^ all-against-all association tests (each calculated for up to 50,000 individuals). Our implementation of *cis*-QTL mapping with permutations to estimate the empirical false discovery rate was >250x times faster than the current state-of-the-art implementation (FastQTL^8^; Fig 1d). Likewise, *trans*-QTL mapping (i.e., 500 billion regressions) took less than 10 minutes, a ∼200x increase in speed compared to running on a CPU (Fig. 1c & Supplementary Fig. 1b). Our current implementation does not scale linearly as a function of samples (Supplementary Fig. 1c) due to limitations in data transfer from the memory of the CPU to the GPU, rather than computational capacity; we leave this additional optimization for future work (Fig. 1e, Supplementary Fig. 1c). We used data from the V6p and V7 releases of GTEx^12^ generated using Matrix eQTL^7^ and FastQTL^8^, respectively, to demonstrate the reproducibility of our implementation (Fig. 1f & Supplemental Figure 1d).

In addition to the savings in computation time, implementation in TensorFlow or PyTorch also results in significant cost savings -- at the time of writing, GPU compute time cost ∼$0.50-$0.75/hour on multiple cloud platforms compared to ∼$0.01-0.05/hour for CPUs, thus the same analyses were ∼5-10 fold cheaper on GPUs.

## Conclusions

In summary, implementation of many commonly used methods in genomics using new GPU-compatible libraries can vastly increase speed and reduce costs compared to CPU-based approaches. Indeed, by simply re-implementing current methods we were able to achieve an order of magnitude higher increase in speed than may be achieved through sophisticated approximations for optimizing runtimes on CPUs^13, 14^. Our findings indicate that the scale of computations made possible with GPUs will enable investigation of previously unanswerable hypotheses involving more complex models, larger datasets, and more accurate empirical measurements. For example, our GPU implementation enables computation of empirical p-values for *trans*-QTL, which is cost-prohibitive on CPUs. Similarly, our results show that GPU-based approaches will enable scaling of single-cell analysis methods to millions of cells. Given the availability of libraries that obviate the need for specialized GPU programming, we anticipate a transition to GPU-based computing for a wide range of computational genomics methods.

## Software Availability

All software is available on Github and implemented in python using open source libraries. TensorQTL: https://github.com/broadinstitute/tensorqtlSignatureAnalyzer-GPU: https://github.com/broadinstitute/SignatureAnalyzer-GPU

## Author Contributions

A.T.W., and F.A. outlined and planned development. A.T.W., F.A., G.G., E.M.V prepared and reviewed manuscript and figures. F.A. and A.T.W developed tensorQTL. A.T.W., N.H., J.K. and S.G. developed SignatureAnalyzer-GPU. F.A. performed and planned benchmarking analysis for tensorQTL. A.T.W., N.H., S.A., and J.K. performed and planned benchmarking analysis for SignatureAnalyzer-GPU. All authors reviewed and edited the manuscript.

## Figure Legends

**Supplementary Figure 1:**
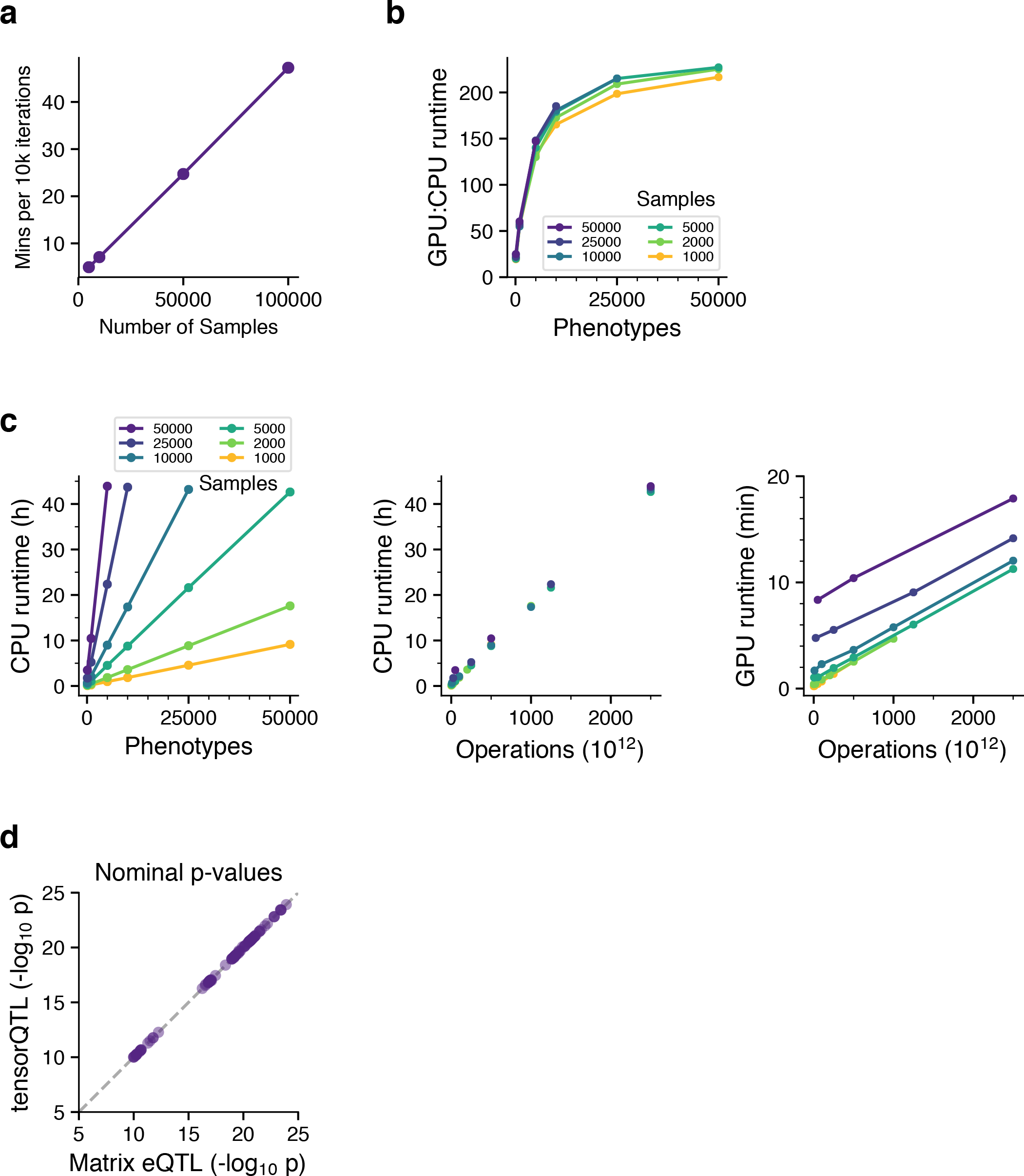
GPU performance scaling of SignatureAnalyzer-GPU and tensorQTL. a) SignatureAnalyzer-GPU runtime scales linearly as a function of number of samples. b) GPU-to-CPU runtime ratio for tensorQTL, across the indicated phenotype and sample sizes, for 10^7^ common variants. The ratio is non-constant due to data load and CPU-to-GPU memory input/ouput times (“i/o”) that are more limiting for large sample sizes (number of individuals). c) CPU runtime of tensorQTL for the indicated range of sample and phenotype sizes (left panel). CPU runtimes scale linearly demonstrated by the collapse of the compute time when measured as a function of number of operations (approximated as phenotypes x samples x variants; middle panel), whereas GPU runtimes do not show this collapse for large sample sizes due to i/o limitations (right panel). d) Nominal significant *trans*-eQTL p-values from the V6p TEx release^12^ replicated using tensorQTL.

## References

1. Bycroft, C. et al. The UK Biobank resource with deep phenotyping and genomic data. Nature 562, 203–209 (2018).

2. McArt, D. G. et al. cudaMap: a GPU accelerated program for gene expression connectivity mapping. BMC Bioinformatics 14, 305 (2013).

3. Mejía-Roa, E. et al. NMF-mGPU: non-negative matrix factorization on multi-GPU systems. BMC Bioinformatics 16, 43 (2015).

4. Schatz, M. C., Trapnell, C., Delcher, A. L. & Varshney, A. High-throughput sequence alignment using Graphics Processing Units. BMC Bioinformatics 8, 474 (2007).

5. Abadi, M. et al. TensorFlow: Large-Scale Machine Learning on Heterogeneous Distributed Systems.

6. Paszke, A., et al. Automatic differentiation in PyTorch. (2017).

7. Shabalin, A. A. Matrix eQTL: ultra fast eQTL analysis via large matrix operations. Bioinformatics 28, 1353–8 (2012).

8. Ongen, H., Buil, A., Brown, A. A., Dermitzakis, E. T. & Delaneau, O. Fast and efficient QTL mapper for thousands of molecular phenotypes. Bioinformatics 32, 1479–1485 (2016).

9. Kim, J. et al. Somatic ERCC2 mutations are associated with a distinct genomic signature in urothelial tumors. Nat. Genet. 48, 600–606 (2016).

10. Alexandrov, L. et al. The Repertoire of Mutational Signatures in Human Cancer. bioRxiv 322859 (2018). doi:10.1101/322859

11. Haradhvala, N. J. et al. Distinct mutational signatures characterize concurrent loss of polymerase proofreading and mismatch repair. Nat. Commun. 9, 1746 (2018).

12. GTEx Consortium. Genetic effects on gene expression across human tissues. Nature 550, 204–213 (2017).

13. Loh, P.-R., Kichaev, G., Gazal, S., Schoech, A. P. & Price, A. L. Mixed-model association for biobank-scale datasets. Nat. Genet. 50, 906–908 (2018).

14. Zhou, W. et al. Efficiently controlling for case-control imbalance and sample relatedness in large-scale genetic association studies. Nat. Genet. 50, 1335–1341 (2018).

